# Coral reef carbonate budgets and ecological drivers in the naturally high temperature and high alkalinity environment of the Red Sea

**DOI:** 10.1101/203885

**Authors:** Anna Roik, Till Röthig, Claudia Pogoreutz, Christian R. Voolstra

## Abstract

The coral structural framework is crucial for maintaining reef ecosystem function and services. In the central Red Sea, a naturally high alkalinity is beneficial to reef growth, but rising water temperatures impair the calcification capacity of reef-building organisms. However, it is currently unknown how beneficial and detrimental factors affect the balance between calcification and erosion, and thereby the overall growth of the reef framework. To provide insight into present-day carbonate budgets and reef growth dynamics in the central Red Sea, we measured *in situ* net-accretion and net-erosion rates (G_net_) by deployment of limestone blocks and estimated census-based carbonate budgets (G_budget_) in four reef sites along a cross-shelf gradient (25 km). We assessed abiotic variables (i.e., temperature, inorganic nutrients, and carbonate system variables) and biotic drivers (i.e., calcifier and bioeroder abundances). On average, total alkalinity A_T_ (2346 - 2431 μmol kg^−1^), aragonite saturation state (4.5 - 5.2 Ω_a_), and pCO_2_ (283 -315 μatm) were close to estimates of pre-industrial global ocean surface waters. Despite these calcification-favorable carbonate system conditions, G_net_ and G_budget_ encompassed positive (offshore) and negative net-production (midshore-lagoon and exposed nearshore site) estimates. Notably, G_budget_ maxima were lower compared to reef growth from pristine Indian Ocean sites. Yet, a comparison with historical data from the northern Red Sea suggests that overall reef growth in the Red Sea has likely remained similar since 1995. When assessing sites across the shelf gradient, A_T_ correlated well with reef growth rates (ρ = 0.89), while temperature was a weaker, negative correlate (ρ = −0.71). Further, A_T_ explained about 65 % of G_budget_ in a best fitting distance-based linear model. Interestingly, parrotfish abundances added up to 82% of explained variation, further substantiating recent studies highlighting the importance of parrotfish to reef ecosystem function. Our study provides a baseline that will be particularly useful in assessing future trajectories of reef growth capacities in the Red Sea under continuous ocean warming and acidification.

## Introduction

Coral reef growth is limited to warm, aragonite-saturated, and oligotrophic tropical oceans and is pivotal for coral reef functioning (Buddemeier, 1997; Kleypas et al., 1999). The coral reef framework not only maintains a remarkable biodiversity, but also provides highly valuable ecosystem services that include food supply and coastal protection, among others (Moberg and Folke, 1999; Reaka-Kudla, 1997). Biogenic calcification, erosion, and dissolution cumulatively contribute to the formation of the reef framework constructed of calcium carbonate (CaCO3, mainly aragonite) (Glynn, 1997; Perry et al., 2008). The balance of carbonate loss and accretion are controlled by abiotic and biotic factors: temperature, properties of carbonate chemistry (e.g. pH, total alkalinity A_T_, and aragonite saturation state Ω_a_), calcifying benthic communities (scleractinian corals and coralline algal crusts), as well as grazing and endolithic bioeroders (e.g. parrotfish, sea urchins, and boring sponges) (Glynn and Manzello, 2015; Kleypas et al., 2001).

Positive carbonate budgets (G_budget_) are maintained when reef calcification produces more CaCO_3_ than is being removed, and rely in a great part on the ability of benthic calcifiers to precipitate calcium carbonate from seawater (Ca^2+^ + CO_3_^−^ ↔ CaCO_3_, Tambutté et al. 2011). Calcification rates increase with higher temperature, but have an upper thermal limit (Jokiel and Coles, 1990; Marshall and Clode, 2004). In addition, A_T_ and Ω_a_, are measures for the availability of the carbonate ions in seawater and their tendency to precipitate. Both positively correlate with calcification rates (Marubini et al., 2008; Schneider and Erez, 2006). Today’s oceans are warming, which poses a potential threat to calcifying reef organisms, as high temperatures begin to exceed the thermal optima of calcifying organisms and thereby slowing down calcification (Carricart-Ganivet et al., 2012; Death et al., 2009). Simultaneously, as ocean acidification is commonly manifested in a decrease in ocean’s pH and hence a decrease of Ω_a_ (Orr et al., 2005), calcification becomes energetically more costly (Cai et al., 2016; Cohen and Holcomb, 2009; Waldbusser et al., 2016). Finally, ocean acidification stimulates destructive processes, e.g. the proliferation and erosive activity of endolithic organisms that counteract reef growth (Enochs, 2015; Fang et al., 2013; Tribollet et al., 2009). A low or negative G_budget_ is generally associated with a disturbance of abiotic conditions that naturally have supported coral reef growth over the past millennia, i.e. specific temperature range, nutrient level, pH, and Ω_a_ (Buddemeier, 1997; Kleypas et al., 1999). Today, in most tropical coral reefs, negative G_budget_ are a hallmark of reef degradation due to an increased intensity or frequency of extreme climate events (Eakin, 2001; Schuhmacher et al., 2005) or local human impacts, such as pollution and eutrophication (Chazottes et al., 2002; Edinger et al., 2000).

A census-based G_budget_ approach is a powerful tool to assess reef persistence of the reef framework allowing for a regional and global comparison of coral reef ecosystems (Kennedy et al., 2013; Perry et al., 2012, 2015). In more recent years, G_budget_ studies revealed that coral reef growth in the Caribbean has decreased by 50%, compared to historical mid- to late-Holocene reef growth. Also, 37% of all reefs studied are reported to be in a net-erosional state (Perry et al., 2013). In the Red Sea, coral reefs are exposed to challenging conditions in terms of high temperature and salinity regimes (Kleypas et al., 1999). Despite its high temperature and high salinity conditions (Roik et al., 2016), the Red Sea supports remarkable reef ecosystems with a coral reef framework along its entire coastline (Riegl et al., 2012). But coral core samples indicate that calcification rates have already been declining over the past decades worldwide, at the Great Barrier reef, the Caribbean, and also the central Red Sea. This decline in coral growth is widely attributed to ocean warming (Bak et al., 2009; Cantin et al., 2010; Cooper et al., 2008). In the central and southern Red Sea, present-day data show reduced calcification rates of corals and calcifying crusts when temperatures peak during summer (Roik et al., 2015; Sawall et al., 2015). While increasing temperatures are seemingly stressful and energetically demanding for calcifiers, high A_T_ values (~ 2400 μmol kg^−1^, Metzl et al. 1989) in the Red Sea are putatively beneficial for carbonate accretion (Tambutté et al., 2011).

It is yet unknown how these present-day stressors and positive drivers of reef growth influence the overall reef net-carbonate production in this region. Availability of G_budget_ data for Red Sea coral reefs is poor (Jones et al., 2015). Aside from one early census-based assessment of the G_budget_ for a high-latitude reef in the Gulf of Aqaba (northern Red Sea), which considered both calcification and erosion/dissolution rates (Dullo et al., 1996), the remaining studies report calcification rates only (Cantin et al., 2010; Heiss, 1995; Roik et al., 2015; Sawall and Al-Sofyani, 2015). In this study we therefore set out to assess abiotic and biotic drivers of reef growth, and to determine the G_budget_ for coral reefs of the central Red Sea. First, to reveal the present-day carbonate chemistry in the region, we examined sites along an environmental cross-shelf gradient during winter and summer. Second, we followed the census-based *ReefBudget* approach (Perry et al., 2012) to estimate net carbonate production states (G_budget_) using reef site-specific biotic parameters. To achieve this, we assessed the abundances and calcification rates of the major reef-building coral taxa (*Porites*, *Pocillopora*, and *Acropora*) and calcareous crusts, along with the abundances and erosion rates of external macro bioeroders (parrotfish and sea urchins). Also, we measured net-accretion/erosion rates (G_net_) *in situ* using limestone blocks deployed in the reefs, which additionally capture endolithic erosion rates. Finally, we explore the correlations of potential drivers on G_net_ and the overall G_budgets_ using the abiotic and biotic data. Our study provides insight into reef growth dynamics and a comparative baseline to assess the effects of ongoing environmental change on reef growth in the central Red Sea.

## Materials and Methods

### Study sites and environmental monitoring

Study sites were located in the Saudi Arabian central Red Sea along an environmental cross-shelf gradient, which was previously described in (Roik et al., 2016) and Roik et al. (2015). Briefly, along this gradient dissolved oxygen increases, but chlorophyll-a, turbidity, and sedimentation decrease from nearshore to offshore, and are subject to seasonal variation (Roik et al., 2016). Data for this study were collected at four sites: an offshore forereef at ~25 km distance from the coastline (22° 20.456 N, 38° 51.127 E, “Shi’b Nazar”), two midshore sites (forereef and lagoon) at ~10 km distance (22° 15.100 N, 38° 57.386 E, “Al Fahal”), and a nearshore forereef at ~3km distance (22° 13.974 N, 39° 01.760 E, “Inner Fsar”). All sampling stations were located between 7.5 and 9 m depth. In the following, reef sites are referred to as “offshore”, “midshore”, “midshore lagoon”, and “nearshore”, respectively. Abiotic variables were measured in the study sites during two seasons in 2014. Temperature and pH were measured continuously recorded during “winter” (10^th^ February – 6^th^ April 2014) and “summer” (20^th^ June – 20^th^ September 2014) (see Roik et al., 2016). Additionally, for 5 – 6 consecutive weeks during each of the seasons, seawater samples were collected on SCUBA at the stations for the determination of inorganic nutrients and carbonate chemistry: nitrate and nitrite (NO_3_^−^&NO_2_^−^), ammonia (NH_4_^+^), phosphate (PO_4_^3−^), total alkalinity (A_T_), and pH (Table S1).

### Abiotic parameters - Continuous data

Conductivity-Temperature-Depth loggers (CTDs, SBE 16plusV2 SEACAT, RS-232, Sea-Bird Electronics, Bellevue, WA, USA) equipped with pH probes (SBE 18/27, Sea-Bird) were deployed at 0.5 m above the reef to collect times series data of temperature and pH_CTD_ at hourly intervals. Both sensors were factory-calibrated. To control for drift, pH probes were tested before and after deployment using certified standard buffers (pH-7 “38746” and pH-10 “38749”, Fluka Analytics, Sigma-Aldrich, Germany). Data corrections were applied if necessary.

### Abiotic parameters - Seawater samples

Seawater samples were collected on SCUBA at each of the stations using 4 L cubitainers (Table S1). Simultaneously, 60 mL seawater samples were taken over a 0.45 μm syringe filter for A_T_ measurements. Immediately after sampling, the pH of the seawater (pHSWS) was measured in subsamples using a portable pH probe with an integrated temperature sensor (n = 3, precision of ± 0.05 pH units, Orion 4 Star Plus, Thermo Fisher Scientific, MA, USA). Before each sampling day, the probe was calibrated using certified standard buffers (pH-4 “38743,” pH-7 “38746” and pH-10 “38749”, Fluka Analystics, Sigma-Aldrich). Seawater samples for inorganic nutrient analyses and A_T_ measurement were transported on ice in the dark and were processed on the same day. Samples were filtered over GF/F filters (0.7 μm, Whatman, UK) and filtrates were frozen at −20 °C until analysis. The inorganic nutrient content (NO_3_^−^&NO_2_^−^, NH_4_^+^, and PO_4_^3−^) was determined using standard colorimetric tests and a Quick-Chem 8000 AutoAnalyzer (Zellweger Analysis, Inc.). A_T_ samples were analyzed within 2 - 4 h after collection using an automated acidimetric titration system (Titrando 888, Metrohm AG, Switzerland). Gran-type titrations were performed with a 0.01 M HCl certified standard solution (prepared from 0.1 HCl, Fluka Analytics) at a precision of ± 9 μmol kg^−1^.

### Abiotic parameters - Net-accretion/erosion rates in limestone blocks (G_net_)

Net-accretion/erosion rates were assessed using a limestone block “assay”. Blocks (100 × 100 × 21 mm, ρ = 2.3 kg L^−1^, n = 4) were weighed before and after deployment on the reefs, where they were exposed to the natural processes of calcification and erosion. Before weighing (Mettler Toledo XS2002S, readability = 10 mg), blocks were autoclaved and dried for a week in a climate chamber at 40°C (BINDER, Tuttlingen, Germany). Blocks were deployed for 6 months (September 2012 - March 2013), and for 12 months (June 2013 - June 2014) at six sites, including the four reef sites and the offshore and nearshore back reefs, and for 30 months (January 2013 - June 2015) in the four reef sites. Upon recovery, blocks were treated with 10 % bleach for 24 - 36 h to remove organic material. G_net_ were expressed as normalized differences of pre-deployment and post-deployment weights (kg m^−2^ yr^−1^).

### Biotic parameters - Benthic community composition

To assess coral reef benthic calcifier and bioeroder communities as input data for the carbonate budgets, we conducted *in situ* surveys on SCUBA along the cross shelf gradient at each of our study sites.The community composition and coverage of coral reef calcifying groups across was assessed during both sampling seasons on SCUBA. We surveyed benthic calcifiers and non-calcifiers and categorized them as follows: % cover total hard coral, % hard coral morphs (branching, encrusting, massive, and platy/foliose), % major reef-building coral families (Poritidae, Acroporidae, and Pocilloporidae), % cover calcareous crusts, % cover algae & sponges). For a detailed description of the benthic surveys please refer to Roik et al. (2015). In addition, benthic rugosity was assessed using the *Tape and Chain Method* (Perry et al., 2012).

### Biotic parameters - Bioeroder populations along the cross shelf gradient

We surveyed the populations for the two main groups of coral reef framework bioerorders, the parrotfishes (Scaridae) (Bellwood, 1995; Bruggemann et al., 1996) and sea urchins (Echinoidea) (Bak, 1994). Surveys were conducted on SCUBA in stationary plots and line transects respectively per site (n = 6 each). For details on the field surveys and data treatment for biomass conversion, refer to the supplementary materials (Text S1).

### Biotic parameters - Reef carbonate budgets (G_budget_)

Reef carbonate budgets (G_budget_, kg m^−1^ yr^−1^) were estimated following the census-based *ReefBudget* approach (Perry et al., 2012) adjusted for the central Red Sea. Site-specific benthic calcification rates (G_benthos_, kg m^−1^ yr^−1^), net-accretion/erosion rates of hard substrate (G_netbenthos_, kg m^−1^ yr^−1^), and erosion rates of crucial macro bioeroders such as sea urchins (E_echino_, kg m^−1^ yr^−1^) and parrotfishes (E_parrot_, kg m^−1^ yr^−1^) were incorporated in the G_budget_ estimates (Fig. 1). A detailed account of calculations is provided in the supplementary materials (Text S1, Equation box S1-3).

**Figure 1.**
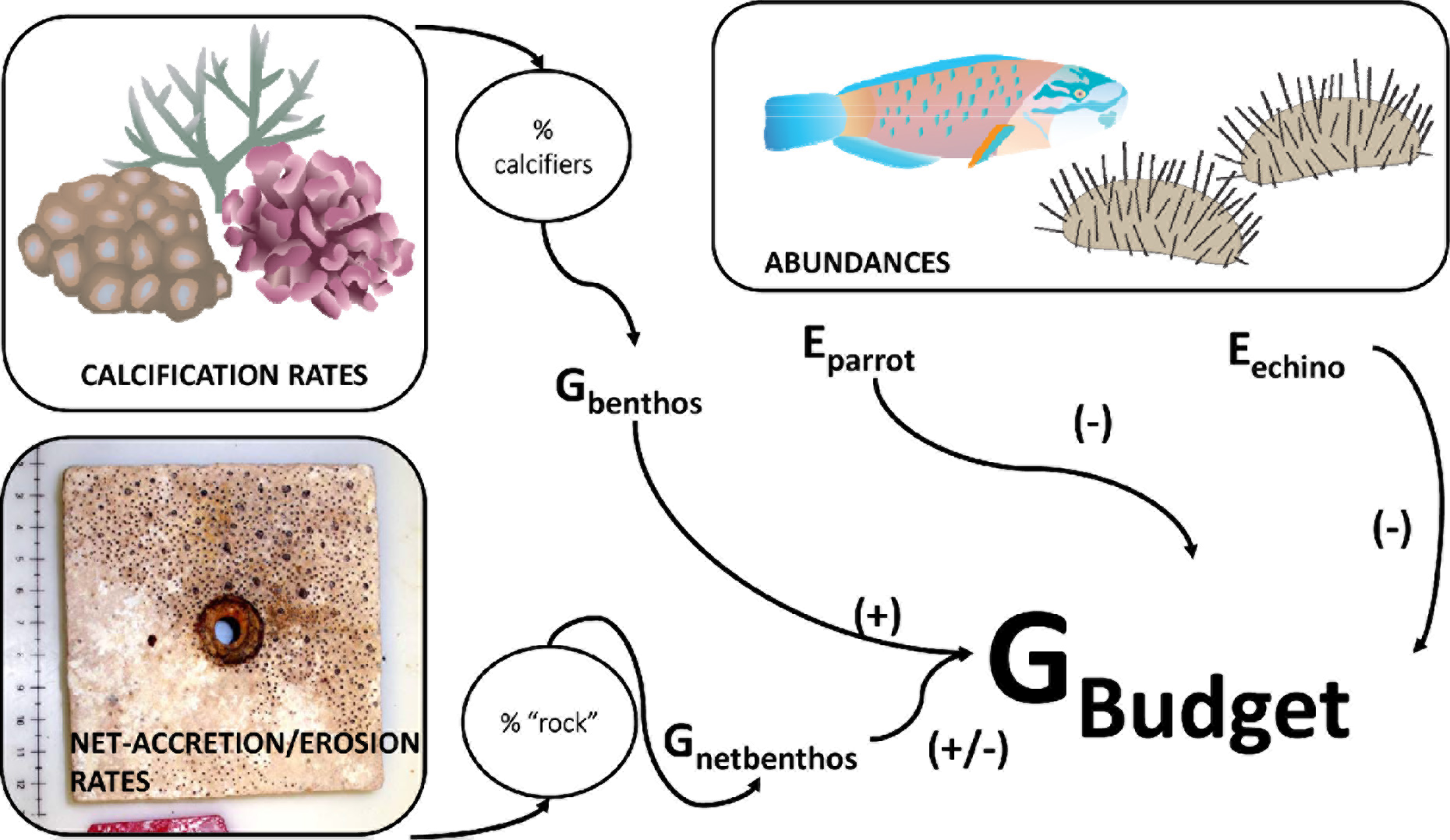
Schematic overview of the census based *ReefBudget* carbonate budget (G_budget_) approach (adapted from Perry et al., 2012). Values and equations that were used are available as Supplementary Materials. G_benthos_ = benthic community calcification rate, G_netbenthos_ = net-accretion/erosion rate of bare reef substrate, E_parrot_ = parrotfish erosion rate, E_echino_ = echinoid (sea urchin) erosion rate, G_budget_ = carbonate budget of a reef. Images from www.ian.umces.edu.

### Statistical analyses - Abiotic parameters

Continuous temperature and pH data were summarized as daily means, daily standard deviations (SD), and daily minima/maxima. Diel profiles were plotted per reef and season including smoothing polynomial regression lines fitted by *geom*_*smooth* in R package *ggplot2* (LOESS, span = 0.1). Data were additionally visualized in histograms using the function *stat bin,* as implemented in the R package *ggplot2* (R Core Team, 2013; Wickham and Chang, 2015). Univariate 2-factorial permutational ANOVAs (PERMANOVAs, Primer-E V6) were used to test the factors “reef” (nearshore, midshore lagoon, midshore, and offshore) and “season” (winter and summer). PERMANOVAS were performed on Euclidian resemblance matrices calculated from log_2_ (x+1) transformed data (Anderson et al., 2008) and were based on 999 permutations of residuals under a reduced model and type II partial sums of squares. Within each significant factor, pair-wise post-hoc tests followed.

Using A_T_, pH_SWS_ (seawater samples), salinity, and temperature (from CTD), carbonate chemistry parameters were calculated using the R package *seacarb* (Gattuso et al., 2015). Carbonic acid dissociation constants were employed as recommended in Dickson et al. (2007): K1 & K2 (Lueker et al., 2000), Kf (Perez and Fraga, 1987), and Ks (Dickson, 1990). Then, inorganic nutrients (NO_3_^−^&NO_2_^−^, NH_4_^+^, and PO_4_^3−^) and carbonate system variables (pH_SWS_, A_T_, C_T_, Ω_a_, HCO_3_^−^, CO_3_^2−^) were evaluated using multivariate PERMANOVAs followed by principal coordinate ordination (PCO) according to the continuous data test design. Multivariate 2-factorial PERMANOVAs on Euclidian resemblance matrices created from normalized data were run under same specifications as above. Next, univariate 2-factorial ANOVAs were employed to evaluate the parameters separately under the same test design. For this, inorganic nutrients and carbonate chemistry parameters were transformed (log_2_ (x+1): pH_SWS_, A_T_, Ω_a_; square-root: C_T_, CO_3_^2−^; box-cox: HCO_3−_) to meet the assumptions of normality and homoscedascity.

### Statistical analyses - Net-accretion/erosion rates and carbonate budgets

G_net_ data were tested for effects of the factors “reef” (nearshore, midshore, and offshore), “site exposure” (fore- and backreef), and “deployment time” (6, 12, and 30 months). Because of the incomplete design due to missing the nearshore and offshore backreef sites in the 30-months deployment, a univariate 3-factorial PERMANOVA was conducted using Euclidian distance matrix 999 permutations of residuals under a reduced model and type II partial sum of squares. G_budget_ were tested for statistical differences between the four “reef sites” (nearshore, midshore lagoon, midshore, and offshore) using a 1-factorial ANOVA, after box-cox transforming the data to meet the assumptions. In parallel, biotic variables were tested using a 1-factorial ANOVA for square-root transformed G_benthos_, and non-parametric Kruskal-Wallis tests for non-transformed G_netbenthos_, E_echino_, and E_parrot_. Tukey’s HSD post-hoc tests followed where applicable.

### Statistical analyses - Abiotic-biotic correlations

To evaluate the relationship of abiotic and biotic predictors of G_net_ and G_budget_, a multivariate statistics approach was applied. Distance-based linear models (DistLM) were performed including biotic and abiotic predictor variables (Primer-E V6). Models were tested for (a) G_net_ and (b) G_budget_ data. G_net_ encompassed data of pooled 12- and 30-months measurements from four reef sites (nearshore, midshore lagoon, midshore, and offshore). Predictor variables in (a) were reef growth relevant abiotic parameters, comprising means and SDs from continuous measurements of temperature and pH_CTD_ per reef site, and the means of inorganic nutrients (NO_3_^−^ &NO_2_^−^, NH_4_^+^, and PO_4_^3^) and carbonate chemistry parameters (A_T_, C_T_, Ω_a_, HCO_3−_, and CO_3_^2^) (Table 1). Biotic variables that can potentially influence G_net_ on limestone blocks were added to the models, i.e. parrotfish abundances and percentage cover (%) of calcareous crusts, both derived from the reef surveys. In (b), the same predictor variables were employed as for (a), but biotic predictors were extended with additional variables available from reef surveys i.e. % cover total hard coral, % hard coral morphs (branching, encrusting, massive, and platy/foliose), % major reef-building coral families (Poritidae, Acroporidae, and Pocilloporidae), % cover calcareous crusts, % cover algae & sponges, benthic rugosity, and abundances of sea urchins and parrotfish. Prior to DistLM, some predictor variables (i.e. sea urchin and parrotfish abundances, % platy/foliose corals, and % Poritidae) were log_10_ (x+1) transformed to improve the symmetry in their distributions following (Anderson et al., 2008). Both DistLM routines were performed on Euclidian resemblance matrices, implementing the step-wise forward procedure with 9999 permutations and adjusted R^2^ criterion. Additionally, Spearman rank correlation coefficients were obtained for the response variables and their predictors.

**Table 1.** Abiotic parameters relevant for reef growth at coral reef sites along a cross-shelf gradient in the central Red Sea. Temperature (Temp) and pH were continuously measured using *in situ* probes (CTDs). Weekly collected seawater samples were used for the determination of inorganic nutrient concentrations, i.e. nitrate and nitrite (NO_3_^−^&NO_2_^−^), ammonia (NH_4_^+^), and phosphate (PO_4_^3−^). Carbonate chemistry parameters were measured as total alkalinity (A_T_) and pH in the same samples and used to calculate the carbonate ion concentration (CO_3_^2−^), aragonite saturation state (Ω_a_), total inorganic carbon (C_T_), bicarbonate ion (HCO_3_^−^), and partial pressure of carbon dioxide (pCO_2_).

|  | Continuous data |  | Seawater samples |  |  |  |  |  |  |  |  |  |
| --- | --- | --- | --- | --- | --- | --- | --- | --- | --- | --- | --- | --- |
| Site / Season | Temp | pH<sub>CTD</sub> | $\text{NO}_3^-$ & $\text{NO}_2^-$ | $\text{NH}_4^+$ | $\text{PO}_4^{3-}$ | pH<sub>SWS</sub> | $A_T$ | $\text{HCO}_3^{*-}$ | $\text{CO}_3^{2-*}$ | $\Omega_a^*$ | $C_T^*$ | $\text{pCO}_2^*$ |
| | °C | | $\mu\text{mol L}^{-1}$ | $\mu\text{mol L}^{-1}$ | $\mu\text{mol L}^{-1}$ | | $\mu\text{mol L}^{-1}$ | $\mu\text{mol kg}^{-1}$ | $\mu\text{mol kg}^{-1}$ | | $\mu\text{mol kg}^{-1}$ | $\mu\text{atm}$ |
| Avg. winter | 26.0<br>(0.6) | 8.13<br>(0.19) | 0.36<br>(0.25) | 0.35<br>(0.2) | 0.07<br>(0.02) | 8.16<br>(0.02) | 2423<br>(18) | 1683<br>(24) | 299<br>(7) | 4.62<br>(0.12) | 1990<br>(21) | 292<br>(14) |
| Avg. summer | 31.0<br>(0.7) | 8.15<br>(0.19) | 0.61<br>(0.25) | 0.5<br>(0.22) | 0.03<br>(0.02) | 8.13<br>(0.03) | 2369<br>(38) | 1588<br>(41) | 314<br>(17) | 4.95<br>(0.28) | 1910<br>(36) | 303<br>(25) |
| Nearshore exposed / winter | 26.1<br>(0.7) | 8.25<br>(0.27) | 0.31<br>(0.17) | 0.34<br>(0.19) | 0.06<br>(0.02) | 8.15<br>(0.02) | 2414<br>(21) | 1678<br>(36) | 297<br>(6) | 4.6<br>(0.11) | 1983<br>(31) | 295<br>(16) |
| Nearshore exposed / summer | 31.5<br>(0.6) | 8.31<br>(0.12) | 0.6<br>(0.28) | 0.42<br>(0.17) | 0.02<br>(0.01) | 8.11<br>(0.03) | 2346<br>(31) | 1576<br>(37) | 309<br>(14) | 4.88<br>(0.23) | 1892<br>(32) | 311<br>(22) |
| Midshore sheltered / winter | 25.9<br>(0.6) | 8.00<br>(0.17) | 0.41<br>(0.31) | 0.46<br>(0.28) | 0.07<br>(0.02) | 8.15<br>(0.02) | 2421<br>(21) | 1695<br>(19) | 294<br>(10) | 4.54<br>(0.14) | 1997<br>(16) | 301<br>(17) |
| Midshore sheltered / summer | 30.9<br>(0.6) | 8.09<br>(0.22) | 0.68<br>(0.26) | 0.58<br>(0.28) | 0.04<br>(0.03) | 8.11<br>(0.03) | 2365<br>(40) | 1603<br>(47) | 306<br>(18) | 4.83<br>(0.29) | 1917<br>(40) | 315<br>(31) |
| Midshore exposed / winter | 26.1<br>(0.5) | 8.15<br>(0.07) | 0.32<br>(0.3) | 0.34<br>(0.19) | 0.06<br>(0.02) | 8.17<br>(0.02) | 2431<br>(14) | 1679<br>(27) | 304<br>(8) | 4.7<br>(0.13) | 1991<br>(21) | 286<br>(15) |
| Midshore exposed / summer | 30.7<br>(0.7) | 8.16<br>(0.09) | 0.58<br>(0.26) | 0.5<br>(0.26) | 0.02<br>(0.01) | 8.13<br>(0.02) | 2373<br>(38) | 1594<br>(37) | 313<br>(15) | 4.93<br>(0.24) | 1915<br>(34) | 300<br>(16) |
| Offshore exposed / winter | 26.0<br>(0.4) | 8.10<br>(0.05) | 0.4<br>(0.24) | 0.26<br>(0.15) | 0.08<br>(0.02) | 8.16<br>(0.01) | 2425<br>(20) | 1679<br>(18) | 302<br>(2) | 4.66<br>(0.03) | 1988<br>(19) | 285<br>(5) |
| Offshore exposed / summer | 30.8<br>(0.7) | 8.12<br>(0.08) | 0.59<br>(0.25) | 0.51<br>(0.18) | 0.03<br>(0.03) | 8.15<br>(0.02) | 2393<br>(35) | 1579<br>(47) | 328<br>(16) | 5.17<br>(0.26) | 1914<br>(39) | 283<br>(19) |

All values as mean (SD); pH<sub>CTD</sub> = pH from CTD; pH<sub>SWS</sub> = pH from seawater samples<sup>#</sup> day-time measurements (7:00 – 19:00); \* parameters calculated using *seacarb* implemented in R (Gattuso et al., 2015; R Core Team, 2013)

## Results

### Abiotic parameters relevant for reef growth - Temperature and pH

The seasonal mean temperature varied between 26.0 ± 0.6°C in winter and 30.9 ± 0.7°C in summer across all reefs (Table 1). The difference across the shelf was on the average ~0.4°C (Table S9). The nearshore and midshore reef experienced the lowest (both ~26°C in winter), and the nearshore reef the highest mean temperatures (31.5 ± 0.6°C in summer). Seasonal and spatial differences in all temperature data (daily means, daily SDs, daily minima and maxima) were significant (Fig. 2 A-B, Table S10). Compared to all other sites, the nearshore reef experienced significantly higher daily maxima during summer (“daily max.”, *p* = 0.01, Fig. 2 B, Table S9), and significantly lower minima during winter (*p* < 0.01, see also Table S10).

**Figure 2.**
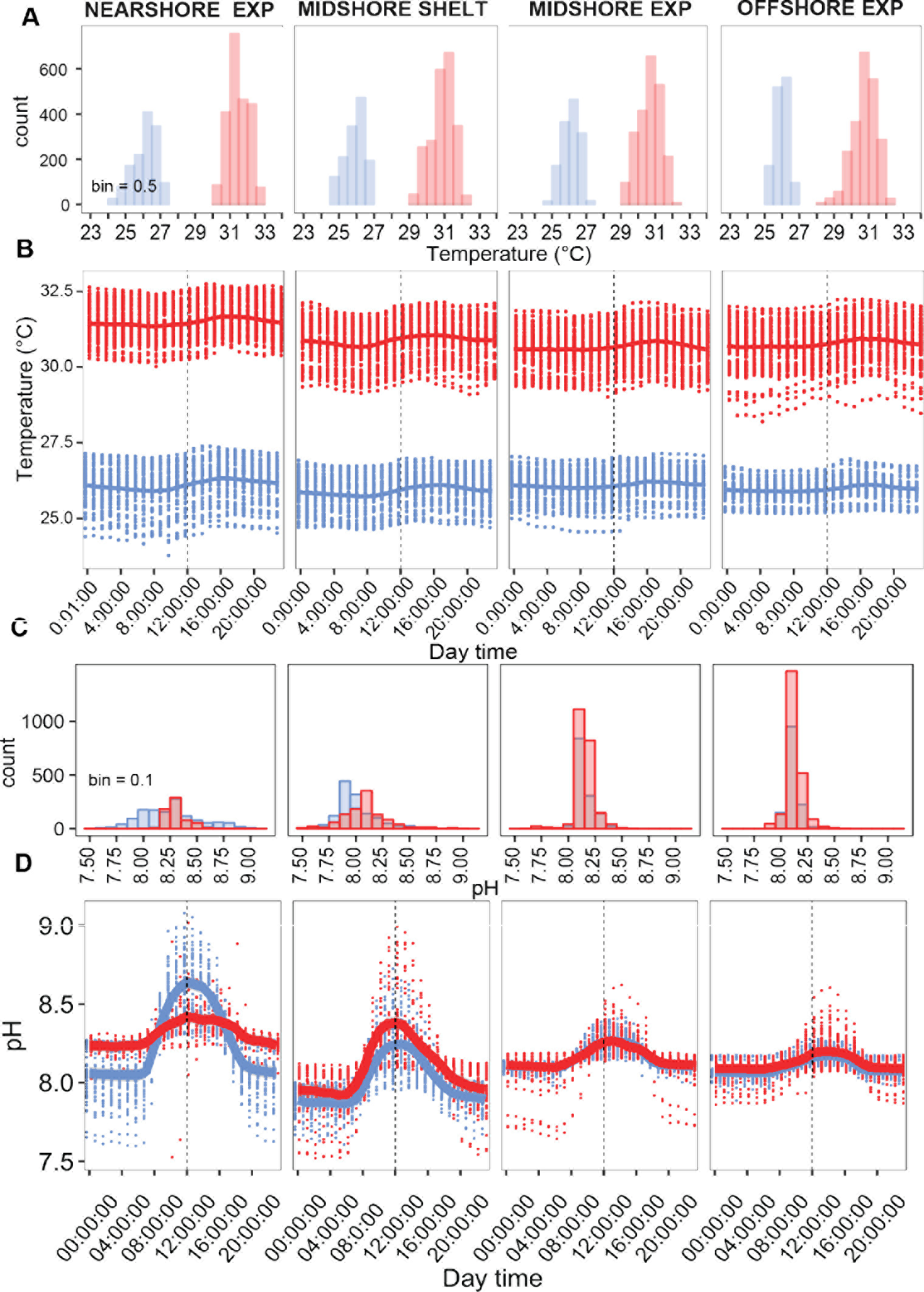
Seasonal temperature and pH regimes on coral reefs along a cross-shelf gradient in the central Red Sea. Continuous data of temperature (A - B) and pH_CTD_ (C - D) collected during winter (blue) and summer (red) at 0.5 m above the reef are presented in histograms (A, C) and diel profiles (B, D). Data points per reef site in winter comprise n = 1287 - 1344, and n = 1099 - 2231 in summer (nearshore summer n = 644). Diel profiles show raw data points and local polynomial regression lines (LOESS, span = 0.1). A dotted vertical line marks the midday time. EXP = exposed, SHELT = sheltered, univar. = univariate, n.s. = not significant, SD = standard deviation, min = minimum, max = maximum

Across all reef sites, seasonal means for pH were 8.13 ± 0.19 in winter and 8.15 ± 0.13 in summer (Table 1). Lowest seasonal means were recorded on the midshore lagoon with 8.00 ± 0.17 in winter and 8.09 ± 0.22 in summer, and highest in the nearshore reef (8.25 ± 0.27 in winter and 8.31 ± 0.12 in summer). pH was intermediate on the exposed midshore and offshore (8.10 ± 0.05 to 8.16 ± 0.09). Overall, continuous pH data showed that spatial differences were more pronounced (with a mean difference between site averages of 0.15 pH units, Table S9), compared to minor effect of seasonality (with a mean difference between seasonal averages of 0. 02 pH units). All daily-pH variables differed between reef sites (*p* < 0.01, Table S10, Fig. 2 C-D). Daily-pH SDs and maxima were significantly different between the seasons (*p* < 0.01 and *p* < 0.05, respectively). On all sites, pH followed a diel pattern with peak values around noon (12:00 h).

### Abiotic parameters relevant for reef growth - Inorganic nutrients and carbonate chemistry

Inorganic nutrients and carbonate chemistry showed a major variation between the seasons (both *p* < 0.001, Table S10, Fig. S1). Specifically, NO_3_^−^&NO_2_^−^ and NH_4_^+^ levels almost doubled in winter (0.36 ± 0.25 μmol L^−1^ and 0.35 ± 0.20 μmol L^−1^) compared to summer (0.61 ± 0.25 μmol L^−1^ and 0.50 ± 0.22 μmol L^−1^). In contrast, PO_4_^3−^ was higher in winter than in summer (0.07 ± 0.02 vs. 0.03 ± 0.02 μmol L^−1^, Table 1, Table S10 and Fig. 3 A). Highest inorganic nutrient contents were measured in the midshore lagoon with up to 0.68 μmol NO_3_^−^&NO_2_^−^ L^−1^, 0.58 μmol NH_4_^+^ L^−1^ in summer, and 0.07 μmol PO_4_^3−^ L^−1^ in winter, but PO_4_^3−^ was also highe on the offshore reef during winter (0.08 μmol PO_4_^3−^L^−1^).

**Figure 3.**
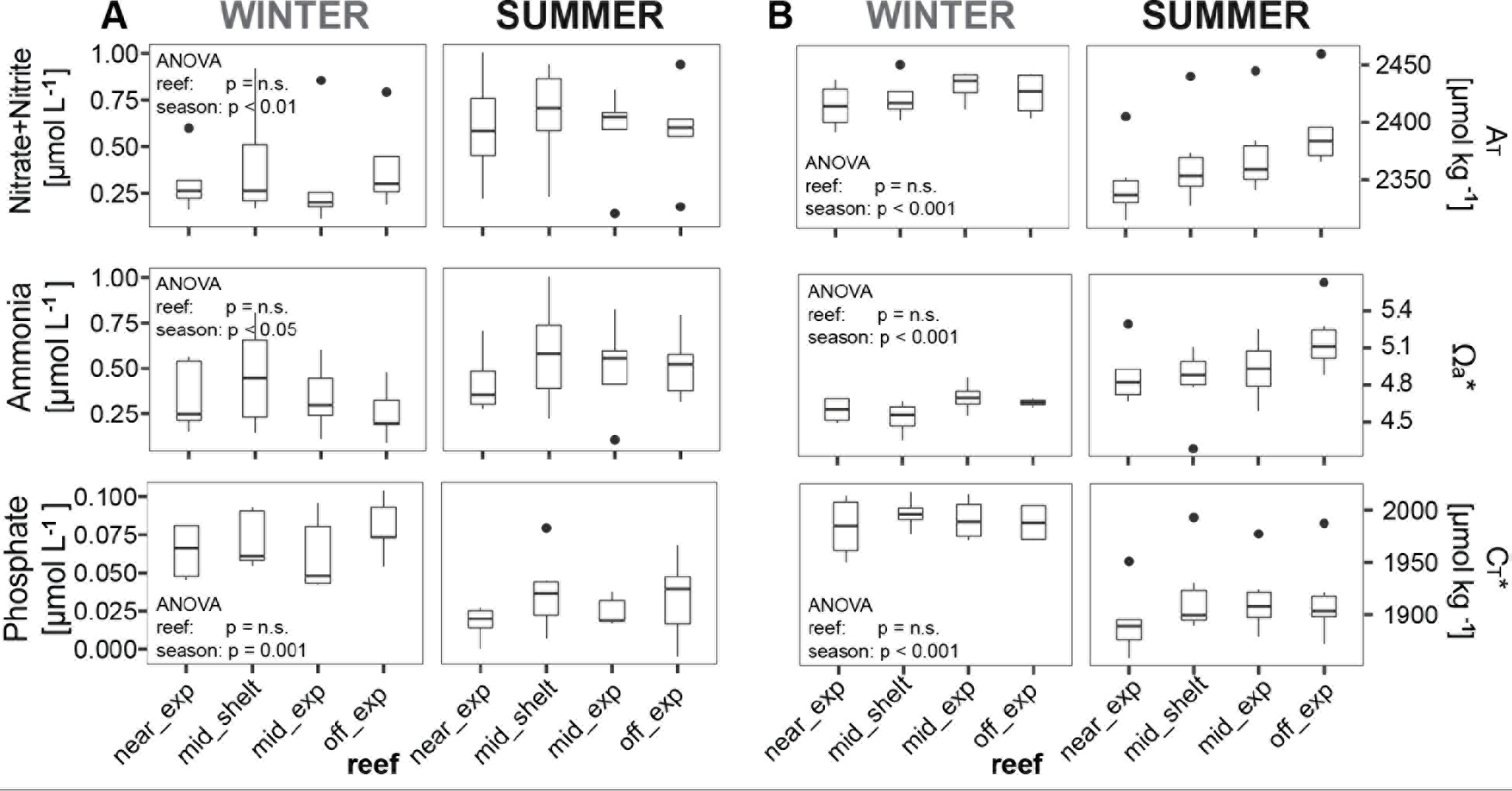
Inorganic nutrients and carbonate system conditions across reef sites and seasons in the central Red Sea. Boxplots illustrate the differences of seawater parameters between the reefs within each season (box: 1st and 3rd quartiles, whiskers: 1.5-fold inter-quartile range, points: data beyond this range). A_T_ = total alkalinity, Ω_a_ = aragonite saturation state, C_T_ = total inorganic carbon; off = offshore, mid = midshore, near = nearshore, exp = exposed forereef, shelt = sheltered lagoon, n.s. = not significant

Carbonate chemistry analysis show overall elevated A_T_, C_T_ and HCO_3_^−^ concentrations in winter (2423 ± 18 μmol A_T_ L^−1^, 1990 ± 21 μmol C_T_ L^−1^, and 1683 ± 24 μmol HCO_3_^−^ L^−1^) compared to summer (2369 ± 38 A_T_ μmol L^−1^, 1910 ± 36 μmol C_T_ L^−1^, and 1588 ± 41 μmol HCO_3_^−^ L^−1^, Table 1, Fig. 3 B, Table S10). Estimates of pCO_2_ in this study ranged 285 - 315 μatm across reef and seasons. C_T_ and HCO_3_^−^ were significantly higher during winter at all sites (*p* < 0.05), while A_T_ was only higher in the nearshore (*p* < 0.05), remaining at similar levels in the other sites. Conversely, Ω_a_ and CO_3_^2−^ were overall reduced during winter (winter: 4.62 ± 0.12 Ω_a_ and 299 ± 7 μmol CO_3_^2−^ L^−1^; summer: 4.95 ± 0.28 Ω_a_ and 314 ± 17 μmol CO_3_^2−^ L^−1^). Changes in Ω_a_ between the seasons were only found in the offshore site (*p* < 0.05). By trend, A_T_ and Ω_a_ increased from nearshore to offshore with average differences of 32 μmol kg^−1^ and Ω_a_ 0.2 (Table S9).

### Biotic parameters relevant for reef growth - Benthic community composition

A detailed account of benthic community structure in the study sites is outlined in Roik et al. (2015). In brief, a low percentage of live substrate (< 40 %) was characteristic of the sheltered and lagoonal sites. In exposed sites (offshore and midshore) a community of calcifying organisms took up to 48 % of benthos cover on average (hard corals and calcareous crusts). Major reef-building corals were the genera *Acropora*, *Pocillopora*, and *Porites* constituting 32-56 % of the total hard coral cover.

### Biotic parameters relevant for reef growth - Abundances and biomasses of macro bioeroders

A total of 718 parrotfishes and 110 sea urchins were observed in the present study. For sea urchins, mean abundances and biomass estimates of 0.002 ± 0.004 − 0.014 ± 0.006 individuals m^−2^ and 0.05 ± 0.04 − 1.43 ± 0.98 g m^−2^ were observed, respectively (Table S4). Parrotfish mean abundances and biomass estimates ranged from 0.05 ± 0.01 − 0.17 ± 0.60 individuals m^−2^ and 19.54 ± 5.56 − 82.18 ± 46.67 g m^−2^, respectively (Table S6). The highest abundances and biomasses of both parrotfishes and sea urchins were observed at the exposed nearshore site. Abundances and biomasses of these two bioeroding groups decreased towards the exposed midshore site, and then increased again towards the exposed offshore site. The inshore sites along with the exposed midshore site exhibited the largest range of sea urchin size classes (from categories 1 or 2 to the largest size class 5), while at the exposed sites, only the two smallest size classes of sea urchins were recorded. The largest parrotfishes (category 5 parrotfish, i.e., > 45 cm - 69 cm fork length) were observed at the midshore sites and the sheltered offshore site. With the exception of the exposed midshore site, category 1 (5 - 14 cm) parrotfish were commonly observed at all sites. In contrast, no category 6 parrotfish (≥ 70 cm fork length) were observed during the surveys.

### Net-accretion/erosion rates

Cumulative net-accretion/erosion rates G_net_ were measured in assays over 6, 12, 30 months in the reef sites along the cross-shelf gradient. Visible boring traces of endolithic worms or sponges were only found on the surfaces of blocks recovered after 12 and 30 months (Fig. 4 A - H). G_net_ based on the 30-months deployment of blocks ranged between −0.96 and 0.37 kg m^™2^ yr^−1^ (Table 2). G_net_ for 12 and 30-months blocks were negative on the nearshore reef (between −0.96 and −0.2 kg m^−2^ yr^−1^, i.e., net erosion is apparent), near-zero on the midshore reef (0.01 - 0.06 kg m^−2^ yr^−1^, i. e., low net accretion), and positive on the offshore reef (up to 0.37 kg m^−2^ yr^−1^, i.e., high net accretion). Reef sites and deployment times had a significant effect on the variability of G_net_ (Table S11). The rate of accretion/erosion was higher in the measurements over longest deployment period (*p* < 0.001, Figure S2).

**Table 2.** Net-accretion/erosion rates G_net_ in coral reefs along a cross-shelf gradient in the central Red Sea, cumulative over 6, 12, and 30 months. G_net_ (kg m^−2^ yr^−1^) was calculated using weight gain/loss of limestone blocks deployed in the reef sites for 6, 12, and 30 months. Means per reef site and standard deviations in brackets. yr = year

| $G_{\text{net}}$ ($\text{kg m}^{-2} \text{yr}^{-1}$) | Deployment time (months) | | |
| --- | --- | --- | --- |
| Reef site | 6 | 12 | 30 |
| Nearshore sheltered | 0.16 (0.09) | -0.2 (0.35) | - |
| Nearshore exposed | 0.11 (0.07) | -0.61 (0.49) | -0.96 (0.75) |
| Midshore sheltered | 0.13 (0.09) | 0.06 (0.03) | -0.29 (0.12) |
| Midshore exposed | 0.11 (0.16) | 0.01 (0.07) | 0.06 (0.12) |
| Offshore sheltered | 0.03 (0.02) | -0.07 (0.07) | - |
| Offshore exposed | 0.14 (0.11) | 0.08 (0.09) | 0.37 (0.08) |

**Figure 4.**
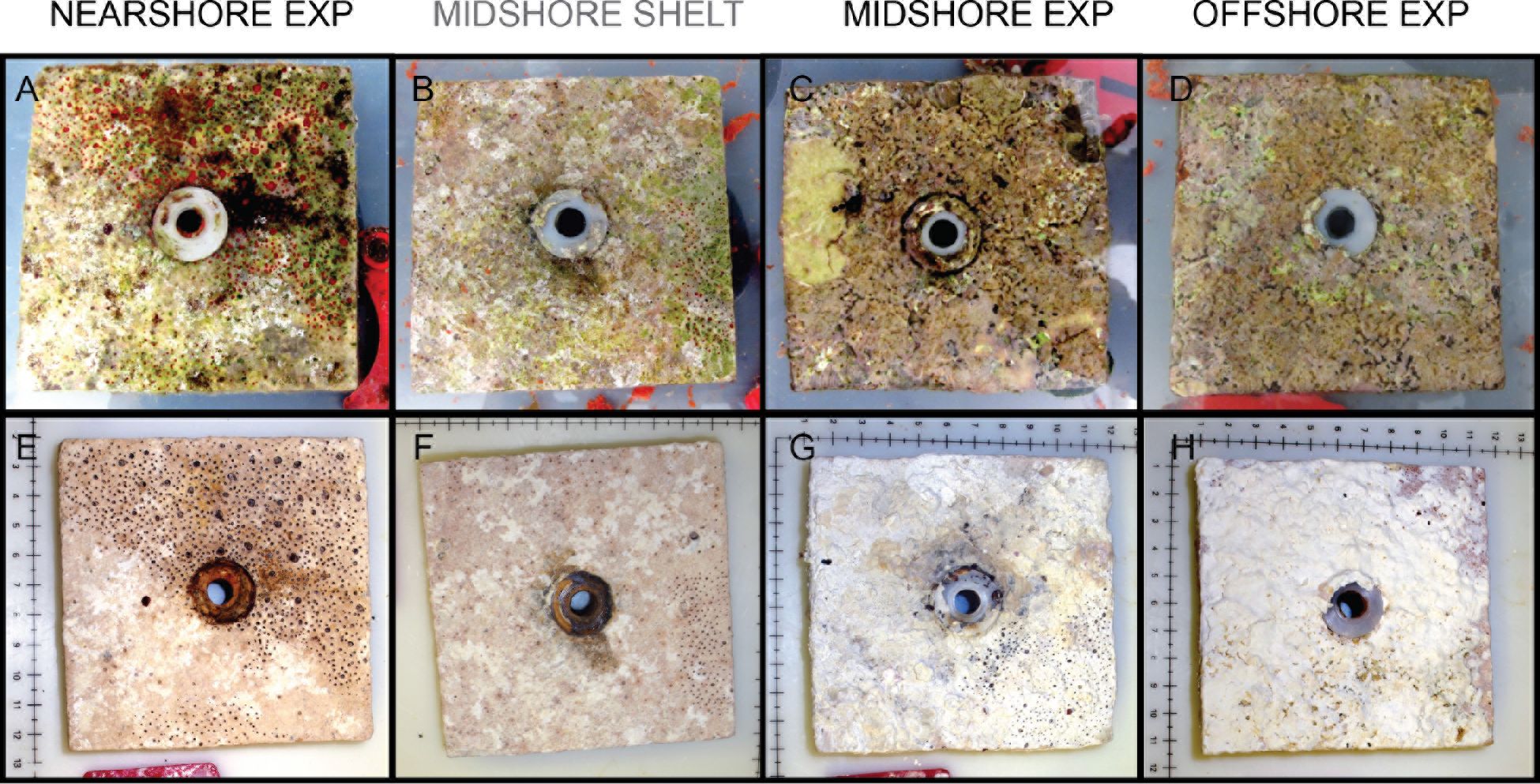
Limestone blocks after 30 months of deployment in the reef sites for measurements of net-accretion/erosion rates G_net_. A - D show freshly collected limestone blocks that were recovered after 30 months, and the same blocks (E - H) after bleaching and drying. Boring holes of endolithic sponges are clearly visible in the nearshore exposed and both midshore reef sites. In the midshore and offshore exposed reefs, blocks were covered with crusts of biogenic carbonate mostly accreted by coralline algae assemblages. EXP = exposed, SHELT = sheltered, scales in E - H in cm.

### Carbonate budgets

The carbonate budget G_budget_, estimated via the *ReefBudget* approach (Perry et al., 2012) and averaged over all sites was 0.65 ± 1.73 kg m^−2^ yr^−1^. This average encompasses values from the negative nearshore budget −1.48 ± 1.75 kg m^−2^ yr^−1^ to the positive offshore budget 2.44 ± 1.03 kg m^−2^ yr^−1^ (Table 3). G_budget_ significantly different between the reef sites (*p* < 0.05, Fig. 5 A), except for budgets in both midshore sites (lagoon and exposed), which were similar. Biotic variables that account for the carbonate budgets also differed by site, in the case of community calcification rates G_benthos_ (*p* < 0.05, Fig. 5 B), and net-accretion/erosion of bare substrate G_netbenthos_ (*p* < 0.001, Fig. 5 C). However, differences of parrot fish and echinoid erosion rates (E_echino_ and E_parrot_) were not significant (Fig. 5 D-E).

**Table 3.** Reef carbonate budget estimates and contributing biotic variables (kg m^−2^ yr^−1^) along a cross-shelf gradient in the central Red Sea. Calcification rates of benthic calcifiers (G_benthos_), net-accretion/erosion rates of reef substrate (G_netbenthos_), and the erosion rates of echinoids and parrotfishes (E_echino_, E_parrot_) contribute to the total carbonate budget (G_budget_) in a reef site. Means per site are shown and standard deviations are in brackets. The last row gives the means and standard deviations across all sites.

| Reef | $G_{\text{budget}}$ | $G_{\text{benthos}}$ | $G_{\text{netbenthos}}$ | $E_{\text{echino}}$ | $E_{\text{parrot}}$ |
| --- | --- | --- | --- | --- | --- |
| Nearshore exposed | -1.477 (1.748) | 0.426 (0.149) | -0.315 (0.129) | -0.228 (0.189) | -1.36 (1.886) |
| Midshore sheltered | 0.598 (0.368) | 1.15 (0.222) | -0.027 (0.014) | -0.187 (0.193) | -0.338 (0.271) |
| Midshore exposed | 1.02 (0.353) | 1.762 (0.242) | 0.009 (0.003) | -0.024 (0.04) | -0.727 (0.307) |
| Offshore exposed | 2.443 (1.033) | 2.812 (0.646) | 0.094 (0.022) | -0.019 (0.003) | -0.444 (0.701) |
| Cross-shelf gradient | 0.646 (1.734) | 1.538 (0.958) | -0.06 (0.168) | -0.114 (0.159) | -0.717 (1.04) |

**Figure 5.**
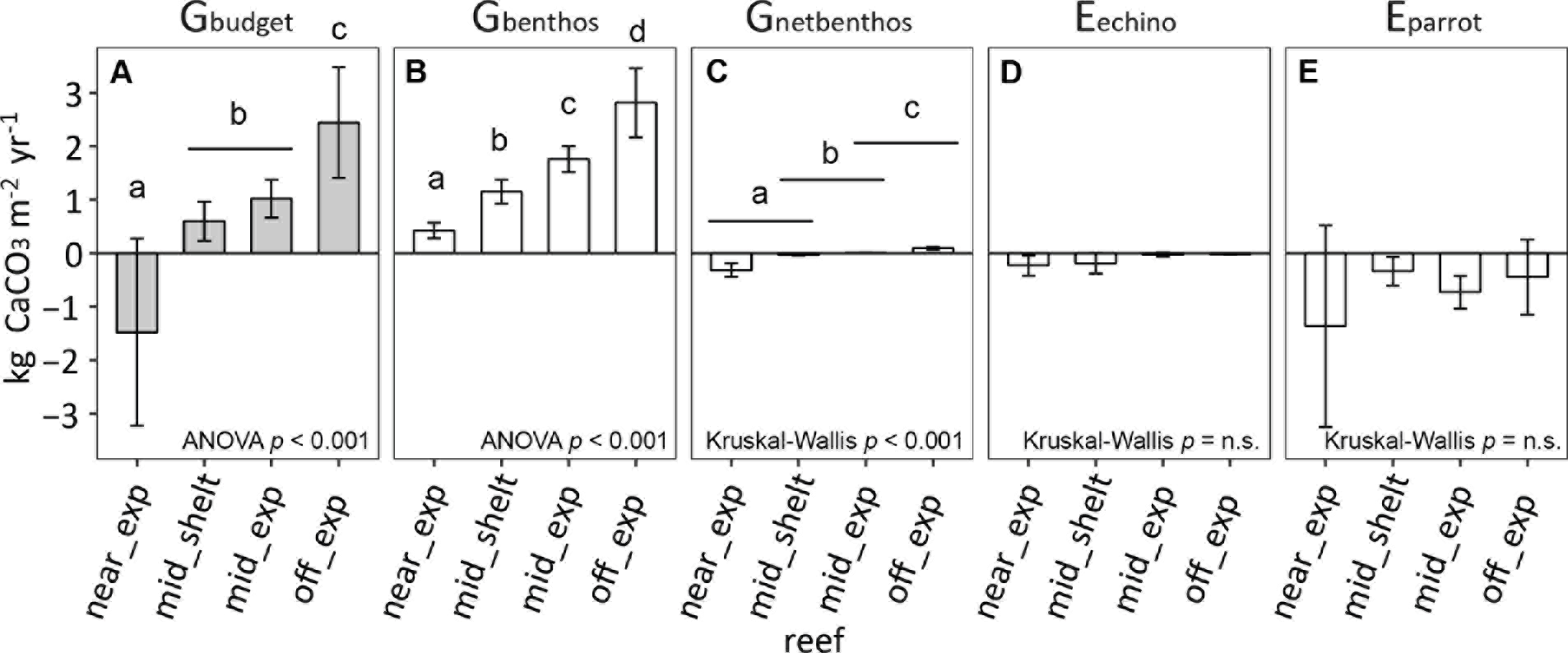
Reef carbonate budget estimates and contributing biotic variables along a crossshelf gradient in the central Red Sea. Benthos accretion (G_benthos_, G_netbenthos_), and the erosion rates of echinoids and parrotfishes (E_echino_, E_parrot_) contribute to the total reef carbonate budget (G_budget_) at each reef site. All data are presented as mean ± standard deviation. (A) G_budget_ and (B - E) biotic variables (G_benthos_, G_netbenthos_, E_echino_, E_parrot_). Letters a - d indicate significant differences between the sites. Near_exp = nearshore exposed, mid_shelt = midshore sheltered (lagoon), mid_exp = midshore exposed, off_exp = offshore exposed.

### Abiotic and biotic drivers related to net-accretion rates and carbonate budgets

Results from correlations and distance based linear models were similar for G_net_ and G_budget_. Temperature means, temperature SDs, and pH SDs were negatively correlated, while A_T_, Ω_a_, CO_3_^2−^, and PO_4_^3−^ were positively correlated with G_net_ (ρ ≥ |0.59|, Table 4). The best model for G_net_ data accounted for 56% (adjusted R^2^) of the total variation. Here, A_T_ alone explained 54% of the data and was the only statistically valid predictor of two abiotic variables in the model (the second being Ω_a_ accounting for only 2% more, Table 5). Negative correlates of G_budget_ were temperature means, temperature SDs, and pH SDs, while A_T_, Ω_a_, CO_3_^2−^, and PO_4_^3−^ were positives (ρ ≥ |0.59|, Table 4). Among biotic variables, % total hard coral and calcareous crusts were positively correlated. The best model for G_budget_ fitted six biotic and abiotic predictors and explained total variation of 87% (adjusted R^2^). Again, A_T_ was the major predictor explaining 65% alone. The biotic variable ‘parrotfish abundance’ added up to 85% of explained variation. Both variables were significant in the test. The remaining four predictors included in the model (NO_3_^−^&NO_2_^−^, % cover total hard coral, % encrusting coral, % of total hard coral, and % Acroporidae) were non-significant and of minute relevance (altogether contributing only 3%, Table 5).

**Table 4.** Coefficients from Spearman rank order correlations for predictor variables vs. G_net_ and G_budget_. 12 abiotic and 13 biotic variables were correlated with G_net_ (= net-accretion/erosion rates of limestone blocks) and G_budget_ (= carbonate budget estimates). Biotic variables encompassed the abundances of bioeroders (echinoids, parrotfishes), and 11 relevant benthic categories (%-cover). Correlates only shown, when Spearman’s correlation coefficient ρ ≥ |0.59|.

| | $G_{\text{net}}$ | | $G_{\text{budget}}$ | |
| --- | --- | --- | --- | --- |
| | $\rho$ | $p$ | $\rho$ | $p$ |
| **<u>Abiotic variables</u>** |  |  |  |  |
| Temperature(ctd) mean | -0.65 | < 0.05 | -0.71 | < 0.05 |
| Temperature(ctd) SD | -0.65 | < 0.05 | -0.71 | < 0.05 |
| pH(ctd) SD | -0.59 | < 0.05 | -0.65 | < 0.05 |
| $A_T$ mean | 0.80 | < 0.05 | 0.89 | < 0.05 |
| $\Omega_a$ mean | 0.59 | < 0.05 | 0.65 | < 0.05 |
| $\text{CO}_3^{2-}$ mean | 0.59 | < 0.05 | 0.65 | < 0.05 |
| $\text{PO}_4^{3-}$ mean | 0.65 | < 0.05 | 0.71 | < 0.05 |
| **<u>Biotic variables</u>** |  |  |  |  |
| Rugosity | - | - | 0.67 | < 0.05 |
| % total hard coral | - | - | 0.70 | < 0.05 |
| % calcareous crusts | 0.59 | < 0.05 | 0.69 | < 0.05 |

**Table 5.**
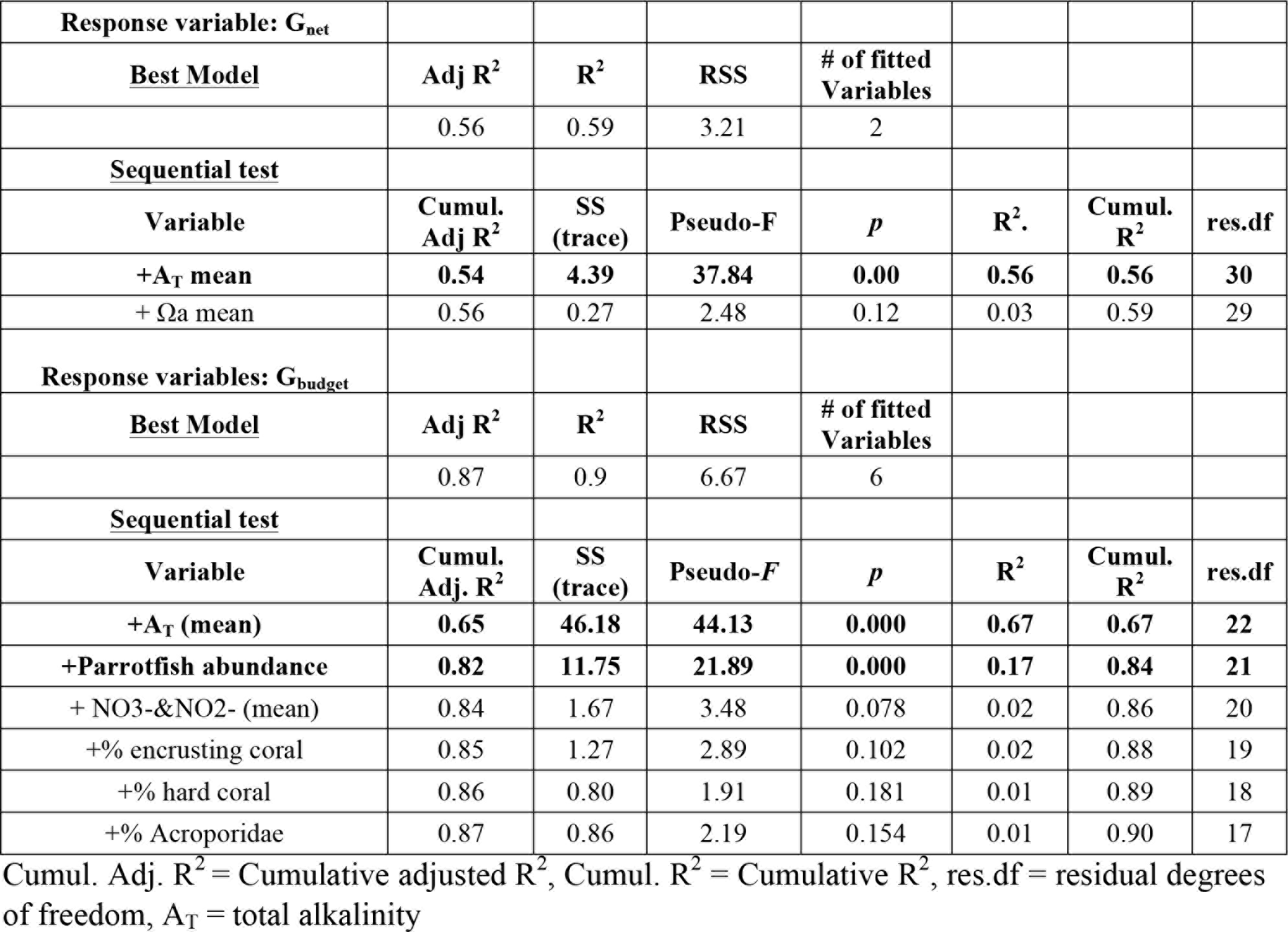
Distance based linear models (DistLM) and sequential tests. Response variables were G_net_ (net-accretion/erosion rates of limestone blocks) and G_budget_ (reef carbonate budget estimates). Predictor variables were 12 abiotic variables, bioeroder abundances (one variable for G_net_, two for G_budget_), and %-cover of relevant benthic categories (one for G_net_, 11 for G_budget_). Significant predictors in **bold**.

| Response variable: $G_{\text{net}}$ | | | | | | | |
| --- | --- | --- | --- | --- | --- | --- | --- |
| <u>Best Model</u> | Adj $R^2$ | $R^2$ | RSS | # of fitted Variables | | | |
|  | 0.56 | 0.59 | 3.21 | 2 |  |  |  |
| <u>Sequential test</u> |  |  |  |  |  |  |  |
| Variable | Cumul. Adj $R^2$ | SS (trace) | Pseudo-F | $p$ | $R^2$ | Cumul. $R^2$ | res.df |
| +$A_T$ mean | **0.54** | **4.39** | **37.84** | **0.00** | **0.56** | **0.56** | **30** |
| + $\Omega_a$ mean | 0.56 | 0.27 | 2.48 | 0.12 | 0.03 | 0.59 | 29 |
| Response variables: $G_{\text{budget}}$ | | | | | | | |
| <u>Best Model</u> | Adj $R^2$ | $R^2$ | RSS | # of fitted Variables | | | |
|  | 0.87 | 0.9 | 6.67 | 6 |  |  |  |
| <u>Sequential test</u> |  |  |  |  |  |  |  |
| Variable | Cumul. Adj. $R^2$ | SS (trace) | Pseudo-$F$ | $p$ | $R^2$ | Cumul. $R^2$ | res.df |
| +$A_T$ (mean) | **0.65** | **46.18** | **44.13** | **0.000** | **0.67** | **0.67** | **22** |
| +Parrotfish abundance | **0.82** | **11.75** | **21.89** | **0.000** | **0.17** | **0.84** | **21** |
| + $\text{NO}_3\text{-\&NO}_2\text{-}$ (mean) | 0.84 | 1.67 | 3.48 | 0.078 | 0.02 | 0.86 | 20 |
| +% encrusting coral | 0.85 | 1.27 | 2.89 | 0.102 | 0.02 | 0.88 | 19 |
| +% hard coral | 0.86 | 0.80 | 1.91 | 0.181 | 0.01 | 0.89 | 18 |
| +% Acroporidae | 0.87 | 0.86 | 2.19 | 0.154 | 0.01 | 0.90 | 17 |

Cumul. Adj. $R^2$ = Cumulative adjusted $R^2$ , Cumul. $R^2$ = Cumulative $R^2$ , res.df = residual degrees of freedom, $A_T$ = total alkalinity

## Discussion

In this study we present environmental data of abiotic and biotic variables affecting present-day reef growth in the central Red Sea, a geographic region that is governed by unique conditions of high salinity and temperature, and that is impacted by ocean warming (Cantin et al., 2010; Raitsos et al., 2011). To date, reef growth, more specifically the calcification rates of reef-building corals in the Red Sea, have mostly been investigated regarding the effect of high temperature (Cantin et al., 2010; Roik et al., 2015; Sawall et al., 2015). Our study is therefore the first to link calcification data on an ecosystem scale with a range of abiotic and biotic variables (carbonate chemistry and nutrient availability; abundance and activity of main bioeroders, respectively). This is achieved by applying a census-based carbonate budget (G_budget_) approach following (Perry et al., 2012). Our approach integrates the net-accretion/erosion rates (G_net_) of six reef sites along the cross-shelf gradient assessed *in situ* using a limestone block assay. Additionally, our study provides an account of carbonate system conditions of the reefs in the central Red Sea, where such data is sparse. In the following, we discuss central Red Sea reef growth rates, which span from net-erosive states in the nearshore reef, to net-accretion in the midshore and offshore reefs, in the context of the prevailing abiotic and biotic drivers. Finally, we discuss our results in a global and historical context.

### Abiotic factors governing reef growth in the central Red Sea

In this study we characterized the present-day abiotic conditions temperature, carbonate chemistry, and inorganic nutrients in the central Red Sea, which are considered environmental parameters that affect calcification and bioerosion on coral reefs. Notably, carbonate system variables, i.e., A_T_, Ω_a_ and pCO_2_, observed in the study sites were on average closer to global preindustrial values rather than to recent measurements from Pacific or Caribbean coral reef systems (Table 6). First, this comparison implies that Red Sea waters provide a beneficial environment for reef calcifiers that depend on availability of carbonate ions (i.e., high Ω_a_, low pCO_2_). Second, Red Sea waters have a high buffering capacity against ocean acidification and will probably protect reefs from this threat in the near future, while reefs outside the Red Sea might soon reach a critically low Ω_a_ threshold, which is projected to bring calcification to a halt (Manzello et al., 2008; Yeakel et al., 2015).

**Table 6.**
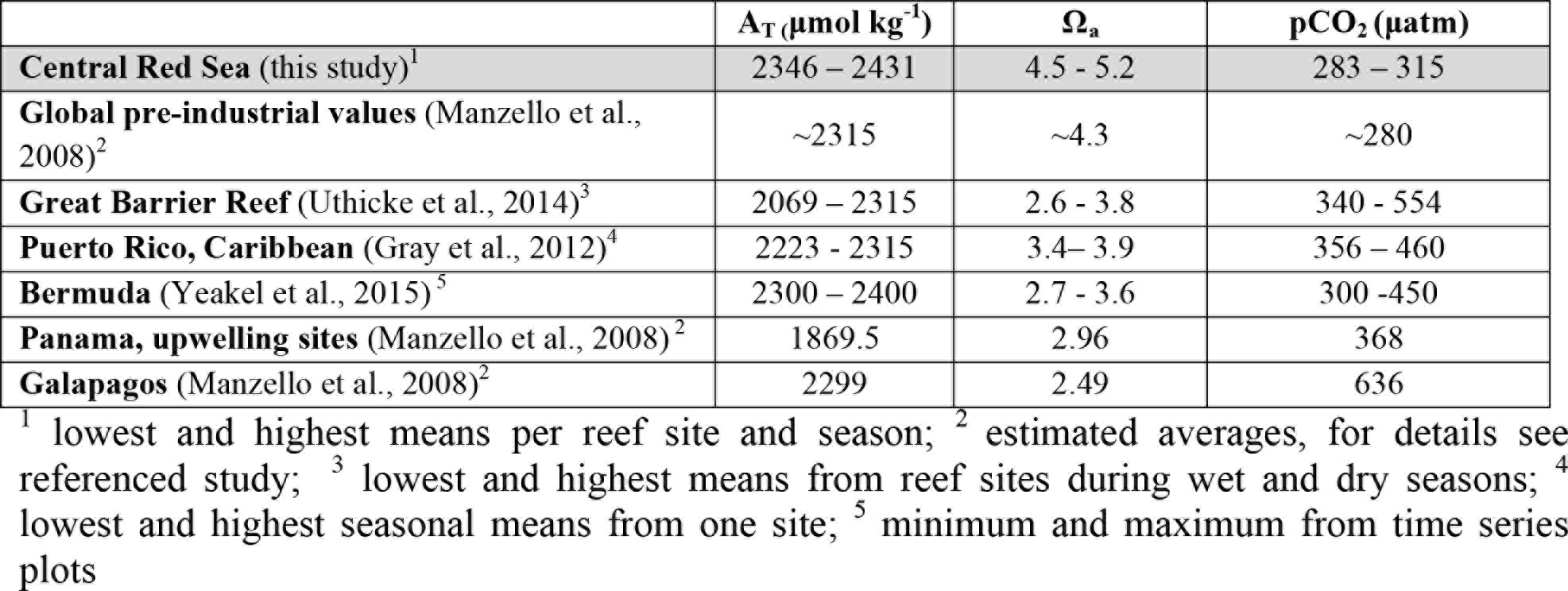
Global comparison of the carbonate system for coral reefs.

In the present study abiotic conditions varied on temporal (seasonal, diel) and spatial scales (cross-shelf gradient, exposed/sheltered reef sites). The observation of the seasonality of the carbonate system was similar to what we know about the high latitude reefs of the GoA, where A_T_ and C_T_ decrease while Ω_a_ increase during summer (Silverman et al., 2007a). While seasonal dynamics of inorganic nutrient concentrations have been shown by remote sensing data for the entire Red Sea basin (Raitsos et al., 2013), the present study demonstrates such dynamics on local reef scale, which again was similar to seasonality in the northern reefs of the GoA (Bednarz et al., 2015). Noteworthy for our study sites was a PO_4_^3−^ enrichment during the winter.

Our continuous recordings show diel fluctuations of temperature and pH that are particularly strong in the nearshore and midshore lagoon sites. Diel pH fluctuation is the consequence of benthic biotic processes, i.e., calcification, dissolution, and respiration/photosynthesis that influence the balance of DIC species through removal, contribution, or exchange of molecules such as CO_3_^2−^, CO_2_, and O_2_, also referred to as biotic feedback (Bates et al., 2010; Silverman et al., 2007a; Zundelevich et al., 2007). The diel pH was more stable in the exposed midshore and offshore site reflecting a weaker biotic feedback, which was likely buffered by the higher rates of reef water mixing with the open sea water (Roik et al., 2016). Diel pH ranges measured on various coral reefs across the globe extend from ~7.80 to ~8.20 pH (Albright et al., 2013; Silbiger et al., 2014) and can span even larger ranges of up to ~1.40 pH units (Hofmann et al., 2011). Such diel pH variations suggest potential co-fluctuation of A_T_ and Ω_a_, which can impact calcification rates at daily time scales. The nearshore and midshore lagoonal reefs from the present study offer suitable study sites to further investigate theses small-scale processes, which to date remain poorly studied.

The difference between nearshore to offshore A_T_ and Ω_a_ in our study area was on average at a range of 32 μmol A_T_ kg^−1^ and 0.2 Ω_a_, respectively (Table S9). Similar cross-shelf differences are reported from e.g. reefs in Bermuda (20 - 40 μmol A_T_ kg^−1^ (Bates et al., 2010). Also in this case, water circulation patterns may explain these spatial gradients. The offshore and midshore reefs receive currents from the Red Sea basin (Roik et al., 2016), which supplies A_T_ saturated open sea water to the reefs. In contrast, the nearshore reef and midshore lagoon are mostly supplied by the boundary current from the south (Roik et al., 2016) travelling along the coastal reef systems which deplete its A_T_.

Indeed, the differences in seawater chemistry between offshore and nearshore reefs were correlated to reef growth processes: the most striking negative correlates were (a) mean temperatures and (b) pH variability, while (c) carbonate system parameters indicative of carbonate ion availability, i.e. A_T_, Ω_a_, CO_3_^2^, and also (d) PO_4_^3−^ concentrations were positively related to reef growth. The negative correlates (a - b) reflect that higher mean temperatures and the impact of strong biotic feedbacks causing pH fluctuations govern nearshore habitats of low reef growth capacity. Previously, pH fluctuation on the micro-habitat scale has been shown to have a significant impact on accretion and erosion dynamics on coral reefs (Silbiger et al., 2014). Potentially these conditions physiologically challenge reef-building organisms exerting a negative effect on the reef growth rate. The positive effect of higher A_T_, Ω_a_, CO_3_^2−^ on the calcification process is well established in laboratory experiments and on reef communities *in situ* (Langdon et al., 2000, Schneider and Erez, 2006, Silverman et al., 2007b, Bates et al., 2010). A_T_ was indeed the strongest predictor for both G_net_ and G_budget_, alone explaining more than half of the variation in reef growth rates in the present study. Interestingly, our study also identifies PO_4_^3−^ concentration, an essential macronutrient and important source of energy for primary producers and reef calcifiers (Ferrier-Pagès et al., 2016), to be a strong abiotic correlate of reef growth. While an overload of inorganic nutrients can be detrimental for the calcification process (Fabricius, 2005; Tambutté et al., 2011), our results show that in a highly oligotrophic reef system, such as the Red Sea, reef growth might be positively affected by seasonal increases in PO_4_^3−^ levels. Experimental studies have shown that PO_4_^3−^ additions can help maintain coral-algae symbiosis in reef-building corals that suffer from heat-stress (Ezzat et al., 2016). Also, under circumstances phosphorus limitation can increase the stress susceptibility of the coral-algae symbiosis (Pogoreutz et al., 2017; Rädecker et al., 2015; Wiedenmann et al., 2013). In the light of our results, it will be of interest to further study spatio-temporal variation of inorganic nutrient ratios to understand their effects on large-scale and long-term trends of reef growth in the central Red Sea.

### Biotic factors governing reef growth in the central Red Sea

Calcifying benthic communities contribute to carbonate production and are considered the most influential drivers for G_budgets_ on global scale (Franco et al., 2016). Loss of coral cover rapidly gives way to increased bioerosion as the critical force of degradation of the carbonate reef framework. This has become particularly apparent in the Caribbean, where G_budgets_ were reported to shift into negative production states when live hard coral cover dropped below 10% (Perry et al., 2013). Similarly, the relevance of benthic calcifying communities (coral and coralline algae) was highly apparent in the present dataset from the central Red Sea: benthos cover percentage of total hard coral and calcareous crusts constituted the strongest positive correlates for G_budget_, and the latter also for the variable G_net_.

Considerably, the community composition and abundances of bioeroders potentially influence reef carbonate budgets (Alvarez-Filip et al., 2009; Bak, 1994; Bellwood, 1995; Bronstein and Loya, 2014; Bruggemann et al., 1996). Our analyses show that parrotfish erosion was a considerable driving force across our study sites. Parrotfish abundances explained ~20% of G_budget_ data variation, reflecting high parrotfish erosion in the nearshore likely contributing to the net erosion state. Parrotfish abundances and biomasses were lowest at the sheltered midshore site, and both increased towards inshore and offshore. Such differences can be attributed to natural (e.g., species distribution, habitat preferences, reef rugosity) and/or anthropogenically-driven factors (e.g., differential fishing pressure; McClanahan, 1994; McClanahan et al., 1994). Indeed, the Saudi Arabian central Red Sea is subject to decade-long unregulated fishing pressure, which has significantly altered overall reef fish community structures and reduced overall fish biomass compared to less impacted regions in the Red Sea (Kattan et al., 2017). In undisturbed coral reefs, parrotfishes are abundant herbivores with the differential capability to remove algal turfs/macroalgae and/or coral reef framework and therefore a complex implications for coral reef growth (Green and Bellwood, 2009): The ecological role of parrotfish grazing is the regulation of benthic algal growth, supporting the recruitment of reef calcifiers and helping maintain a coral-dominated state (Mumby, 2006). Hence, low parrotfish abundances would primarily reduce erosion pressure on a reef, but secondarily promote phase-shift to non-calcifying organisms, such as fleshy macroalgae (Hughes et al 2007). In the long term, this can cause a decrease in gross carbonate production. Moreover, the overfishing of (parrot)fishes can reduce feeding pressure on bioeroders or their larvae (e.g., sea urchins), resulting in an uncontrolled population increase leading reefs the trajectory of degradation driven by expanding bioeroder populations (Edgar et al., 2010; McClanahan and Shafir, 1990).

### Cross-shelf patterns of net accretion/erosion rates and carbonate budget estimates

Net accretion/erosion rates (G_net_) were measured in limestone block assays, in which blocks were exposed to natural levels of biotic CaCO_3_ accretion (mainly by encrusting calcifiers), endolithic erosion by boring organisms, and surface abrasion by grazing fish along a cross-shelf gradient (Tribollet and Golubic, 2005). The resulting cumulative G_net_ reflects the colonization progress on the limestone blocks (Chazottes et al., 1995; Tribollet and Golubic, 2005). Differences in G_net_ along the cross-shelf gradient however were only observed after an exposure time of greater than 12 months, when well established epilithic and endolithic communities were apparent (Figure 4, Figure S2). Measurements from the 30-months assay reveal a net-erosion state (negative rates) in nearshore and net-accretion state (positive rates) in the offshore reef habitat. On average, G_net_ rates were in a similar range as observed in the reefs of Moorea (−0.49 - 0.63 kg m^−2^ yr^−1^, (Pari et al., 1998). Yet, net-erosion states in the Red Sea did not reach the most extreme erosive scales reported from e.g. Moorea or the Andaman Sea, where lowest values were ~ − 7 to − 4 kg m^−2^ yr^−1^ (Pari et al., 1998; Schmidt and Richter, 2013).

G_budgets_ represent the cumulative contribution of the major biotic drivers of reef growth (G_benthos_, G_netbenthos_, E_echino_ and E_parrot_) for each site (Glynn, 1997; Perry et al., 2012) and resulted in a net-erosive budget in the nearshore reef, low net-accretion (near zero) in the midshore reef, up to a high net-accretion budget in offshore. Increasing across reef sites from nearshore to offshore, G_budgets_ imply that nearshore reefs currently erode with half the speed that the offshore reefs grow, which may be interpreted as the formation of an offshore barrier reef in the central Red Sea (see Figure 5). The cross-shelf dynamics of G_budgets_ and the biotic drivers (G and E) are complex and follow unique patterns that are in parts distinct from what we know from other reef systems. Other than observed in the GBR, where reef growth is reported to be high at inshore reefs (Browne et al., 2013), our nearshore study site, was net-erosive. Also, parrotfish erosion was highest in the nearshore area in the present Red Sea study, whereas lower rates were reported for the inshore reefs in the GBR (Hoey and Bellwood, 2007; Tribollet et al., 2002). On Caribbean islands, parrotfish erosion rates are higher in leeward reefs (that are similar to protected nearshore habitats), but these sites are typically characterized by a high coral cover which drives a positive G_budget_ (Perry et al., 2012, 2014). Unlike in the Red Sea nearshore reef, which had the highest parrotfish erosion, but a negative G_budget_ due to low coral cover. This interregional comparison demonstrates that patterns encountered in one cross-shelf reef system should not necessarily be extrapolated to another system. In conclusion, *in situ* studies will be required for each unique system to understand its dynamics and its responses to environmental change.

### Global and historical perspective on reef growth in the Red Sea

The central Red Sea G_budgets_ are comparable with a majority of reef sites in the Caribbean, eastern and central western Pacific raning from −0.8 to 4.5 kg m^−2^ yr^−1^ (Mallela and Perry, 2007). The highest G_budgets_ from the Red Sea are in the range of average global reef growth. However, Red Sea reefs do not reach highest accretion estimates reported, e.g, from the remote reefs in the Indian Ocean at the Chagos Archipelago which still hold the record with up to 9.8 kg m^−2^ yr^−1^ (Perry et al., 2015).

Due to lacking comparative data, it remains difficult to draw a historical perspective on G_budgets_ in the Red Sea. Among the data available are pelagic and reefal net carbonate accretion rates, estimated using basin-scale historical measurements of A_T_ from 1998 (Steiner et al., 2014). Another data set is the census-based budget approach from a fringing reef in GoA from 1994 - 1996 ((Dullo et al., 1996). Both, the A_T_-based reef accretion estimate from 1998 (0.9 kg m^−2^yr^−1^) and the GoA fringing reef budget from 1994 –1996 (0.7 - 0.9 kg m^−2^yr^−1^), constitute a good match. As well, G_budgets_ assessed in the present study are in accordance with these historical data providing a comparison that supports the notion of stable reef growth rates in the Red Sea over the most recent decades. Additionally, the gross calcification rate of benthic communities (G_benthos_) from our offshore site compares well with the maxima measured in GoA reefs in 1994 (2.7 kg m^−2^ yr^−1^, Heiss, 1995).

Reef growth data from the Red Sea region is sparse, but suggest that coral reef growth might have not changed over the past 20 years despite the ongoing warming trend (Raitsos et al., 2011). However, comparisons of data from the central Red Sea and GoA should be interpreted with caution. Due to the strong latitudinal gradient of temperature and salinity between the central Red Sea and GoA, reef growth dynamics from the two regions may differ and introduce bias in the comparisons. Especially, in the light of a recent study demonstrating that increasing warming rates of sea surface temperatures since 1990 coincided with decreasing coral calcification rates in the central Red Sea (Cantin et al., 2010), it remains to be determined whether this declining calcification capacity has an impact on the overall reef growth. In this context, the data presented in this study will serve as a valuable baseline for comparative future studies in the central Red Sea region. Importantly, these data were collected before the Third Global Bleaching Event, which impacted the region during summer 2015 (Monroe et al., in review) and 2016. Hence, our report will be of great value when assessing potential (long-term) changes in the Red Sea G_budget_ after that significant disturbance.

## Conclusions

The Red Sea represents a geographic region where coral reefs thrive under naturally high temperature, high salinity, and high alkalinity conditions. Baseline data for reef growth from this region are valuable as they provide insight into reef functioning under remarkably variable abiotic conditions that deviate from the global average for coral reefs, and can potentially provide a window into future ocean scenarios. Our data show that carbonate chemistry compares to estimates of preindustrial global ocean surface water, suggesting low susceptibility to ocean acidification for reef calcifiers in the Red Sea. Yet, it remains to be determined how small-scale changes in carbonate chemistry will affect overall reef growth in the central Red Sea under prevailing high temperatures. The offshore reef growth rate from the central Red Sea are comparable to other regions of this world and historical Red Sea data suggest that rates might have not decreased over the past few decades, but rather remained unchanged since 1995 in the Red Sea. Our study shows that carbonate budgets are a powerful tool to track the trajectories of modern-day and future reef states. Our data should provide a valuable baseline and foundation data for evaluating the impact of disturbances, such as the most recent high temperature anomalies on the reef-building communities and the overall reef growth capacity in the central Red Sea.

## Acknowledgements

We thank CMOR for assistance with field operations. This study was supported by funding from King Abdullah University of Science and Technology (KAUST).

